# Comparing the ability of embedding methods on metabolic hypergraphs for capturing taxonomy-based features

**DOI:** 10.1101/2025.07.10.663860

**Authors:** Mattia Cervellini, Blerina Sinaimeri, Catherine Matias, Alessio Martino

## Abstract

**Background:** Metabolic networks are complex systems that describe the biochemical reactions within an organism through pairwise interactions between chemical compounds. While this representation is widely used to study biological function, it fails to capture the full structure of metabolic reactions, many of which involve more than two compounds. Hypergraphs offer a more natural representation, where nodes represent metabolites and hyperedges represent reactions involving multiple participants. Clustering such metabolic hypergraphs can reveal systematic differences among evolutionarily distinct organisms, providing insight into ecological constraints and evolutionary pressures.

**Methods:** In this study, we investigate how different graphs and hypergraphs embedding methods influence their unsupervised clustering, with the goal of capturing taxonomy-based classes. We apply 14 distinct embedding strategies to a large-scale dataset of 8,467 metabolic hypergraphs. Each embedding was followed by hierarchical clustering using a fixed linkage method. To assess performance, we compared the resulting clusters against known taxonomic groupings.

**Results:** Our findings show that the choice of hypergraph embedding has a significant effect on clustering outcomes. Among the tested methods, Bag of Hyperedges with Jaccard distance, Histogram Cosine Kernel, and a Hypergraph Auto-Encoder consistently performed best. We also advocate that the embedding method should be chosen based on the goal of the downstream task.

## 1 Introduction

Metabolic networks describe the full set of biochemical reactions within an organism through pairwise interactions between metabolites. They are central to understanding how organisms grow, survive, and adapt, and have applications in fields ranging from evolutionary biology to biotechnology [1–3]. Because usually reactions involve multiple metabolites, hypergraphs, which naturally model multi-way relationships, offer a more accurate representation than standard graphs [4–6].

It is natural to expect that the topological properties of metabolic networks or hypergraphs reflect systematic differences among evolutionarily distinct groups of organisms (i.e. taxonomy-based classes). These structural patterns, shaped by ecological constraints and evolutionary pressures, may offer insights that go beyond sequence similarity or pathway annotation. In this context, clustering metabolic networks or hypergraphs provides a way to uncover such patterns by grouping species based on their global metabolic organization [7, 8].

To perform clustering on structured data such as graphs or hypergraphs, a common approach is to compute low-dimensional embeddings that capture their topological features. These embeddings allow complex objects to be compared in a common vector space using standard clustering algorithms. In the case of hypergraphs, one widely used strategy is to first convert the hypergraph into a graph by replacing each hyperedge with a clique over its vertices, a process known as *clique expansion* or clique projection; then apply graph embedding algorithms. However, clique expansion is lossy because it does not preserve the full structure of the hypergraph. In most cases, it is not possible to reconstruct the original hypergraph from the clique-expanded graph, even when using the dual representation [9].

An alternative is the *star expansion*, where the hypergraph is transformed into a bipartite graph: one part represents the original nodes, and the other represents hyperedges, with edges connecting each node to the hyperedges it belongs to [10]. While this representation is structurally complete, provided the two parts are clearly distinguished, most downstream methods (e.g., embeddings) treat all nodes in the bipartite graph homogeneously. This uniform treatment ignores the fundamental asymmetry between entities and relationships, often leading to embeddings that fail to capture the higher-order structure of the hypergraph [11, 12]. For these reasons, a variety of hypergraph embedding methods have been proposed in recent years, each aiming to capture different aspects of higher-order structure. It is therefore important to understand how these embeddings differ in their ability to preserve biologically relevant information, particularly in downstream tasks such as clustering.

### Our Contribution

In this paper, we investigate how the choice of embedding influences the unsupervised clustering of metabolic networks modeled as hypergraphs, with the ultimate goal of capturing taxonomy-based classes through the clusters. Specifically, we apply 14 different embedding strategies (some taken from the literature and some proposed by us) to the same large-scale set of metabolic hypergraphs and then cluster the resulting embeddings using a fixed (classical and robust) hierarchical linkage algorithm. By holding the clustering method constant, we ensure that differences in clustering outcomes reflect the expressiveness of the embeddings rather than artifacts of the algorithm. Our goal is to assess which embedding methods and spaces best capture the topological distinctions encoded in the higher-order interactions that align with known biological classifications, providing insight into the comparative utility of current hypergraph representation techniques in real biological settings.

### Related work

We mention that methods relying on metabolic networks (thus pairwise relations) and aiming at recovering taxonomy-based classes appear for instance in [8]. It is known that the problem is inherently difficult: while distinguishing Eukaryotes vs Prokaryotes (C1 level classes below) may be achieved with a good strategy, these authors mention that “Instead, within Prokaryotes, Bacteria and Archaea cannot be clearly distinguished” by their own method, and in fact by most of these approaches. A similar graph-based approach, where nodes correspond to pathways and edges link two pathways whether they share at least one compound, has also been used in [13]. In [14], the authors compare 11 carefully selected organisms (1 Eukaryote, 6 Bacteria, 4 Archaea) in terms of several graph invariants and statistics, showing that the metabolic networks of Archaea are fundamentally different with respect to those of the other two domains. In [15], the authors propose a web tool able to analyze metabolic networks and they evaluate its capabilities in finding a “core reaction graph” (that is, a graph collecting common reactions) at kingdom taxonomy level. Graph-based approaches have also been applied for research questions different from the ones we face in this work (e.g., functional classification and phylogenetic analysis [16–19]). Given the availability of ground-truth taxonomic classification, several other research works employed supervised approaches to predict (that is, classify) the taxonomic classification of an organism starting from its metabolic network. To the best of our knowledge, no such works exist that leverage on hypergraph embedding; conversely, works such as [20–23] employ feature engineered from graph representations or graph embedding to perform classification.

The paper is organized as follows: we first describe our large-scale data in Section 2 and discuss its characteristics as well as the taxonomic classes we aim to recover. Then Section 3 presents the embeddings methods compared in our analysis, as belonging to 3 main classes of methods: Bag of Words (BoW) type methods (Section 3.1), kernels methods (Section 3.2) and Neural Networks (NN) based methods (Section 3.2). A summary of these 14 strategies is given in detail in Table 1. The performance evaluation and quality criteria used in our comparison are introduced in Section 4. Results are presented in detail in Section 5, focusing first on each class of methods (BoW, kernels and NN) and then enlarging the view to select the best performers among all methods (Section 5.4). Finally, conclusions and perspectives are given in Section 6.

**Table 1:** Summary of the 14 embedding strategies stratified by their respective family, type of input representation and size of the resulting embedding space.

| Family | Method | Representation | Size | Reference |
| --- | --- | --- | --- | --- |
| BoW-inspired | BoH w/ Jaccard distance | hypergraph | 3,236 | [20, 56] |
|  | BoH w/ Manhattan distance | hypergraph | 3,236 | [20, 56] |
|  | BoN w/ Betweenness Centrality | clique expansion | 2,834 | Proposed |
|  | BoN w/ Degree Centrality | clique expansion | 2,834 | Proposed |
|  | BoN w/ Node Degree | hypergraph | 2,834 | Proposed |
|  | BoN w/ PageRank | clique expansion | 2,834 | Proposed |
| Kernel-based | Histogram Kernel | hypergraph | - | [28] |
|  | Jaccard Kernel | hypergraph | - | [28] |
|  | Edit Kernel | hypergraph | - | [28] |
|  | Stratified Edit Kernel | hypergraph | - | [28] |
| Neural Networks-based | Graph2Vec | clique expansion | [215–281]<sup>†</sup> | [43] |
|  | HGAE (Complete) | hypergraph | 400 | Proposed |
|  | HGAE (Hyperedges-Only) | hypergraph | 400 | Proposed |
|  | HGAE (One-Hot+Hyperedges) | hypergraph | 400 | Proposed |

<sup>†</sup>Depending on the problem: 215 (C1 and C2), 281 (C3), 272 (C4).

## 2 Data Collection

The dataset includes a total number of 8,467 organisms whose metabolic reactions have been downloaded from the Kyoto Encyclopedia of Genes and Genomes [KEGG database, 24] using the Python libraries Bioservices [25] and BioPython [26]. The set of metabolic reactions of each organism has been parsed as a hyperedge list, collecting in each hyperedge (chemical reaction), the compounds (substrate and products) participating in that reaction. Each node has also been labeled with the corresponding KEGG compound identifier.

Each organism (thus hypergraph) has been lately mapped with the four hierarchical classes-levels provided by the KEGG BRITE taxonomy^1^ [27] that classifies the organism itself at four different levels of granularity across the Linnaeus taxonomy:

1. cellular organization (hereinafter C1)
2. domain/kingdom (hereinafter C2)
3. class (hereinafter C3)
4. genus (hereinafter C4).

Such progressively finer grouping supports both biological interpretation and grouping of metabolic reactions. Specifically:

*C1* level classifies organisms based on their fundamental cellular structure in 2 classes: Eukaryotes (organisms with a nucleus and complex cellular compartmentalization) and Prokaryotes (organisms without a membrane-bound nucleus);

*C2* level indicates the highest-level taxonomic domain to which the organism belongs. It corresponds to one of the 6 major divisions of life: Animals, Archaea, Bacteria, Fungi, Plants and Protists;

*C3* level provides a finer taxonomic grouping below the domain level, typically representing a class, order, or phylum, depending on the organism. This classification helps capturing evolutionary relationships and ecological roles more precisely;

*C4* level corresponds to the genus of the organism, a taxonomic rank grouping species that are closely related and often functionally similar. This classification is particularly useful when analyzing metabolic patterns or pathway conservation across related organisms.

Given the considered levels of taxonomy, a related issue is the non-homogeneous sizes of the classes in each level (C1 to C4), meaning that large classes (i.e, containing many organisms) co-exist with much smaller classes at the same level. This obviously complicates the task of any unsupervised clustering approach.

## 3 Methods: hypergraphs embeddings

In the following, we provide a theoretical overview of all methods exploited to generate embeddings starting from metabolic hypergraphs.

As anticipated in Section 2, in the case of metabolic hypergraphs, nodes represent chemical compounds (or metabolites) and hyperedges represent chemical reactions in which the former are involved. Each of the nodes has its own label which makes it possible to interpret metabolic hypergraphs as node-labelled simplicial complexes where node identifiers belong to a finite set of categorical candidates (i.e., the union of all the compound IDs across all considered metabolic hypergraphs, [28]). This particular aspect of metabolic hypergraphs will be crucial for all the embedding strategies that do follow.

All of the herein presented methods were subsequently fed into the same hierarchical clustering algorithm based on Ward linkage [29]. This choice of an efficient, robust and widely used algorithm was made to ensure fairness of comparison among all of the presented embedding methods. Indeed, by keeping fixed the clustering algorithm that operates on the embedding spaces, we may focus our discussion on the informativeness of each embedding space (that is, how separable are the metabolic hypergraphs once embedded in each Euclidean space). Also, contrarily to k-means, the hierarchical clustering approach provides in one-shot a whole family (hierarchy) of clusterings with different number of clusters, a characteristic that will be useful as we do not have a simple way to select an estimated number of clusters from one hand; and want to compare classifications at different levels (thus with different number of clusters) on the other hand.

### 3.1 Bag of Words Inspired Methods

This family of methods was inspired by a foundational concept of Natural Language Processing (NLP) known as Bag Of Words (BoW). In NLP, the BoW approach represents a document by counting the occurrences of each unique token (i.e., individual words) within the document itself, resulting in a vector whose length equals the number of distinct words in the corpus. The same concept has been adapted to metabolic networks [20] by exploiting the fact that it is possible to create a universal “alphabet” of chemical compounds for all organisms. Additionally, each compound appears only once in each metabolic network. These properties make it so that one could superimpose the set of metabolic reactions of different species and the only difference would be in the number of chemical reactions in which each node takes part (i.e., degree of the node). In accordance with such properties, two types of methods were developed: *Bag of Nodes* and *Bag of Hyperedges*.

#### 3.1.1 Bag of Nodes (BoN)

The BoN approach aims at creating an embedding vector which is the same size as the alphabet of chemical compounds (2,835 in our dataset). The vector may be subsequently filled with various node statistics [30] calculated starting from either the hypergraph itself or its clique projection [31]. Let us recall that the clique projection of a hypergraph can be computed by substituting each hyperedge with a clique of the corresponding size (e.g., hyperedge {A, B, C} is replaced by the edges {A, B},{B, C},{A, C}). It is impossible to recover the original structure from the projection but thanks to this transformation one can calculate almost all nodes statistics typical of graph structures also on hypergraphs (see e.g., [32, 33]). In this context, we rely on the following classical and widely used node statistics:

- Degree;
- Degree Centrality;
- Betweenness Centrality [34];
- Pagerank [35].

One embedding method corresponds to one choice of a node statistic among the 4 listed here. In practice, we computed the first statistic (degree) on the hypergraph, while the other 3 were computed on the clique projection graph.

#### 3.1.2 Bag of Hyperedges (BoH)

The BoH approach is a natural evolution of BoN as it leverages the boundedness of the nodes’ alphabet to create an alphabet of hyperedges (i.e., chemical reactions). The size of such set could theoretically be orders of magnitude greater than the size of the set of nodes, however when this concept is applied to metabolic hypergraphs (and more generally in sparse graphs or hypergraphs) their sizes are comparable. In our context, we have 3,236 different metabolic reactions which thus corresponds to the embedding dimension. The embedding vector in this case is a binary vector *f* (*H*) for each organism or hypergraph *H* where each entry is set to 1 if the organism performs such chemical reaction and to 0 otherwise. Due to the binary and sparse nature of such embedding vectors, the distance metrics chosen to cluster the embeddings were the Jaccard distance [36] and the Manhattan distance. Conversely, for plain BoN, we used the standard Euclidean distance.

### 3.2 Hypergraphs Kernels

Graph kernels are among the most widely used techniques for machine learning tasks on graph structures [37]. A kernel function evaluated between two given objects *i* and *j* can be shown to correspond to the inner product between high-dimensional (or infinite-dimensional) feature maps of the input objects in a Reproducing Kernel Hilbert Space [38–40]. Kernel methods became popular in the late ‘90s and early 00’s in the context of Support Vector Machines (SVMs), as they allowed a maximal-margin linear classifier such as SVMs to solve non-linear pattern classification problems.

The methods presented in this section follow the approach described in [28]. That is, building hypergraph kernels by treating them as simplicial complexes. The following paragraphs provide a brief overview of the four kernel methods employed for clustering metabolic hypergraphs. For further details, the interested reader is referred to the original paper [28].

#### 3.2.1 Histogram Cosine Kernel (HCK)

This method leverages the symbolic histogram technique, which generates explicit embedding by counting the number of meaningful substructures in a structured object (e.g, simplices in a simplicial complex). In our context, this reduces to counting the number of hyperedges (metabolic reactions) in each hypergraph (organism). So the initial embedding process is the same as for the BoH (see Section 3.1.2) with *f* (*H*) denoting the binary vector for hypergraph H indicating which hyperedge is present in that hypergraph.

The Histogram Cosine Kernel (HCK) between any two hypergraphs *H_i_, H_j_*is afterwards computed as the cosine similarity between their respective vector representations, hence:

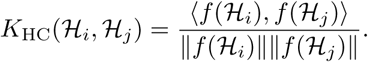

#### 3.2.2 Jaccard Kernel (JK)

The Jaccard Kernel (JK) starts again from the same symbolic histogram as the BoH and the HCK but it is computed as the ratio between the intersection and the union of the two simplicial complexes. Considering *ε*(*H*) as the set of hyperedges of hypergraph H, the kernel in our case is defined as:

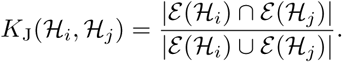

To better understand the links between the different kernels, note that HCK can also be written as

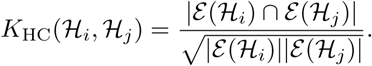

#### 3.2.3 Edit Kernel (EK)

The Edit Kernel (EK) measures the similarity between two hypergraphs considering the number of hyperedges that need to be inserted, removed or substituted in order to transform one hypergraph into the other. Considering *d_L_*(*H_i_, H_j_*) the Levenshtein distance [41] between two hypergraphs, it is possible to convert it to a similarity metric as follows [42]:

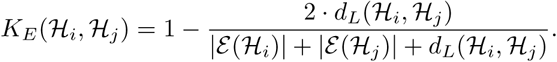

#### 3.2.4 Stratified Edit Kernel (SEK)

The Stratified Edit Kernel (SEK) is a natural evolution of the EK, it avoids the excessive relevance of hyperedges of the most common order (i.e., size). The way it achieves this is by averaging the kernels calculated for each hyperedge size. For each hypergraph *H* and integer *m* ≥ 2, we let H^(*m*)^ denote the sub-hypergraph induced by hyperedges in *H* with size *m* (*m* = 2 corresponding to simple edges, i.e. simple binary relations). Letting *M* denote the largest hyperedge size, the SEK can be defined as:

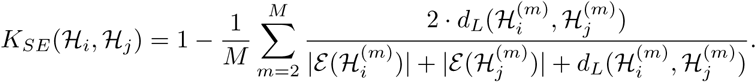

### 3.3 Neural Network-Based Methods

The methods in this section leverage Neural Networks architectures to extract relevant hypergraphs representations.

#### 3.3.1 Graph2Vec

The Graph2Vec approach was first presented in Narayanan et al. in 2017 [43] as an unsupervised graph embedding method inspired its by NLP counterpart, Doc2Vec [44]. Graph2Vec is based on the concept of rooted subgraphs as they represent a high order substructure of graphs as well as being innately non-linear, which conveys more informative representation of graphs [43]. The model works in two main phases: the algorithm first creates rooted subgraphs of node neighborhoods using Weisfeiler-Lehman (WL) relabeling process [45] which natively leverages node labels. This first phase generates a set of rooted subgraphs for the target corpus of graphs; such set is then fed into simple feedforward neural network model commonly known as Skip-Gram model [46] to generate embeddings. In order to accommodate the potentially enormous number of unique rooted subgraphs generated by WL iterations, “negative sampling” is added to the model [see 43, for more details]. This allows training the Skip-Gram model by sampling a small subset of rooted subgraphs as negative examples rather than considering the entire vocabulary.

In order to optimize the hyperparamarers of the Graph2Vec model, Bayesian optimization was applied with the objective of maximizing the Hopkins statistic [i.e., maximizing cluster tendency, 47] of the resulting embeddings dataset. The Hopkins statistic is known to be a fair estimator of randomness in a data set [48]. It is computed by sampling uniformly at random a set W of *m* points from the embedding space, and generating an equal number of uniformly distributed “synthetic” reference points (set *U*). Let [*w*_1_*, …, w_m_*] (resp. [*u*_1_*, …, u_m_*]) be the distances of the points in *W* (resp. in *U*) to their nearest neighbors in the original embeddings dataset. The Hopkins statistic *S* is then defined as:

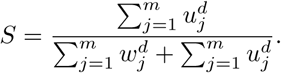

where *d* is the number of dimensions. Values of *S* ≈ 0.5 indicate randomness, while values *S* close to 1 signal strong clustering structure in the dataset [48, 49].

Finally, is it worth mentioning that as Graph2Vec does not natively account for hypergraphs, it was applied to the clique projection of the available metabolic hypergraphs.

#### 3.3.2 Hypergraph Auto-Encoder (HGAE)

The Hypergraph Auto-Encoder (HGAE) method was inspired by the typical structure of Auto-Encoder models, a type of neural network architecture aimed at producing an output as close as possible to the original input it was fed with [50]. Just like most encoder-decoder architectures, the HGAE proposed in this paper passes input information through a bottleneck layer before reconstruction: the process is therefore lossy by design with the aim of retaining the most relevant characteristics of the input, while disregarding noise or less relevant features [51]. The core idea was to train a HGAE model to reproduce the incidence matrix of hypergraphs and then extract the latent representation learned in the bottleneck layer of the model and use it as embedding vector. Considering *n* the number of nodes in a hypergraph and *m* as the number of hyperedges, the incidence matrix can be defined as **H ∈** {0, 1}*^n×m^* such that entry **H***_ij_* = 1 if and only if node *i* takes part in hyperedge *j*, creating a complete representation of the connectivity of the input hypergraph. Hypergraph convolution [52] is used inside the encoder part of the model for message passing among nodes. The HGAE leverages the boundedness of the sets of both nodes and hyperedges, assigning at each distinct element its own dense learnable embedding, it then represents each hypergraph using two matrices: **N** ∈ R*^n×p^* for nodes and **M** ∈ R*^m×p^* for hyperedges, with *p* being the embedding dimension. The decoder part of the model simply computes **Ĥ**∈ R*^n×m^* = **NM***^T^* and model weights (and learnable embeddings) get adjusted after every hypergraph pass considering the reconstruction error between **H** and **Ĥ**. The process described generates node-level embeddings and hyperedge-level embeddings. In order to transform them into hypergraph-level representations, the model considers their average. Among the various versions of HGAE tested, three were included in the paper due to their diverse structures:

- HGAE-Complete
- HGAE-Hyperedges-Only
- HGAE-One-Hot+Hyperedges.

The second and the third variations of HGAE can be interpreted as partials versions of the model structure described in this section. The second variation does not exploit hypergraph convolution during encoding: it leverages only learnable embeddings followed by matrix multiplication (during decoding), the hypergraph level embedding for this version is then computed by considering only the hyperedge embeddings. The third variation, on the other hand, uses convolution but does not involve learnable embeddings for node representations: one-hot encoding is applied for every node label and then message passing is applied before the decoding step. The final embedding in this case considers both nodes and hyperedges.

### 3.4 Summary of compared algorithms

In this section, we have introduced the 14 embedding strategies (summarized in Table 1) whose performances and characteristics will be described in the next section. To foster variety in our comparison, such 14 embedding strategies belong to 3 different families: BoW-inspired, Kernel-based and Neural Network-based (introduced in Sections 3.1, 3.2 and 3.3, respectively).

All BoW-inspired methods yield explicit embedding spaces spanned by *N* × *M* instance matrices, where *N* is the number of hypergraphs in the dataset and *M* is the number of features (that is, number of unique nodes or number of unique hyperedges – see Sections 3.1.1 and 3.1.2, respectively). On the plus side, such explicit embedding spaces foster interpretability as each feature is unambiguously defined and well-characterized (that is, each feature corresponds to either a node or a hyperedge – see Sections 3.1.1 and 3.1.2, respectively). However, they share the negative side of BoW-based modelling in NLP: such matrices can be potentially huge, depending on the size of the input hypergraphs.

Kernel methods provide, as instead, an implicit embedding under the form of pairwise evaluations of a kernel function. In other words, kernel methods yield an *N* × *N* conditionally positive definite matrix scoring, in position (*i, j*) the value of the kernel function evaluated between two given objects *i* and *j*. As kernel matrices endow the pairwise similarities between objects [53, 54], they can directly be used as input (dis)similarity matrices for clustering algorithms such as hierarchical clustering or other distance-based pattern recognition models [55]. Although kernel methods are well-known for their solid mathematical background rooted in statistical learning theory, they lack interpretability due to their implicit embedding.

Lastly, the two Neural Network-based models do yield explicit vectorial embeddings (like BoW-inspired methods), but they are not interpretable (like Kernel-methods and conversely to BoW-inspired methods). Another difference with respect to BoW-inspired methods is that they yield a (usually) smaller embedding space which is compact and dense [43]. However, they learn their embeddings in a fully data-driven, unsupervised manner: latent embeddings from auto-encoders (Section 3.3.2) and Graph2Vec (Section 3.3.1) are not interpretable because they are optimized for reconstruction (the former) or similarity (the latter), not human understanding. They produce entangled representations where each dimension blends multiple abstract features without clear semantic meaning.

## 4 Methods: Evaluation criteria

This section presents the performance criteria used to evaluate the methods.

A first relevant evaluation criteria is the goodness of the entire clustering solutions. The problem at hand does not allow to define an optimal number of clusters for each classification level a-priori, being clustering an unsupervised learning problem, and the main objective was to place organisms belonging to the same class in the same cluster. For these reasons, the possibility of the same class of organisms being spread among various clusters was not regarded as a major concern in this context, as long as most clusters retained a high degree of purity – that is, they contain organisms belonging to the same class. After such considerations, the performance metric of choice for evaluating the goodness of clustering solutions was the *homogeneity*: an entropy-based external cluster valuation measure bounded in the range [0, 1] with 1 being the most desirable score [57].

Formally, let C be the set of classes in the dataset (i.e., coming from the KEGG BRITE taxonomy, see Section 2) and K denote a set of clusters (i.e. obtained through the embeddings). We let *H*(C | K) be the conditional entropy of the classes given the clusters and *H*(C) be the entropy of the classes and respectively defined as:

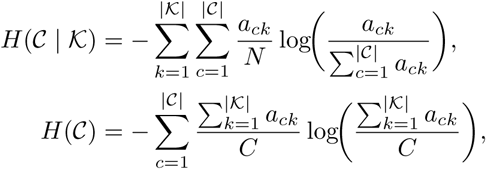

where, in turn, *N* is the number of data points (i.e. the number of organisms or hyper-graphs), *C* is the number of classes (that depends on the level under consideration) and *a_ij_*is the number of data points belonging to the *i*^th^ class lying in the *j*^th^ cluster. The conditional entropy *H*(*C* | *K*) will be close to 0 when the class distribution within each cluster tends to be skewed to a single class. This quantity is compared to the entropy *H*(*C*) which represents the maximum reduction in entropy the clustering information could provide. Finally, the homogeneity *h* of a clustering solution is defined as:

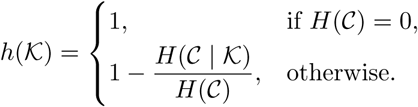

The homogeneity is maximized when each cluster is composed entirely of entities belonging to a same specific class hence, when the conditional entropy of the clusters is 0. Note that when the classes distribution degenerates, *H*(*C*) = 0 and there is a single class, so the clusters will also (by force) be homogeneous. The biggest limitation of homogeneity is that it increases naturally as the number of clusters increase: for this reason it is not helpful in selecting an appropriate number of clusters.

After the overall evaluation of the clustering solutions, a more detailed analysis was conducted on the top-performing methods. This involved examining the resulting clusters to gain deeper insight into how organisms belonging to different classes are spread across the resulting clusters. In this context the performance metric used for each individual cluster was the *purity*, defined as the ratio of elements in a cluster belonging to the majority class in the cluster: its range is (0, 1] and 1 is the most desirable value [58].

Formally, considering K*_k_* the *k*^th^ cluster in a clustering solution K and C = {*c*_1_*, …, c_C_*} the set of *C* true class labels, the purity *p* of a cluster can be defined as follows:

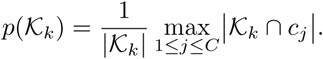

This choice was driven mainly due to its superior intuitiveness compared to homogeneity.

In order to select an appropriate number of clusters for each embedding method at each classification level, it was decided to select the solution presenting the best *silhouette score* [59] by considering a symmetric portion of the dendrogram returned by the hierarchical clustering algorithm centered at the cut point where the number of clusters equals the number of classes for a given classification level or granularity (C1, C2, C3 or C4). Formally, the silhouette score *s*(*x_i_*), for a given data point *x_i_*, is defined as:

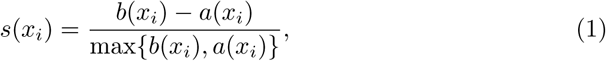

where *a*(*x_i_*) is the average distance between *x_i_*and all the other data points in its own cluster and *b*(*x_i_*) is the nearest-cluster average dissimilarity of which *x_i_* is not a member. Finally, the overall silhouette (i.e., for the whole clustering solution) is taken by averaging each data point’s score:

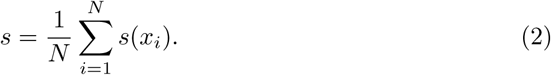

By definition, *s*(*x_i_*) ∈ [–1, +1] (∀*i* = 1*, …, N*) and, by extension, *s* ∈ [–1, +1] as well: the closer to +1, the better the clustering solution.

An important specification to be made is that, due to classifications introduced in Section 2 becoming finer and finer from C1 to C4, the number of classes in each level increases exponentially while the number of organisms for each class decrease drastically. When applying the hierarchical clustering algorithm to the finer classification levels (C3 and C4) it was decided to cut the dataset in order to keep only classes having a significant number of observations according to the following rationale:

- in the case of C3 level, any class containing at least 100 observations was retained: bringing the number of organisms to be clustered to 7, 622 and the number of distinct classes to 18;
- for C4 level, any class containing at least 20 observations was retained: the number of organisms went to 3, 237 and the number of distinct classes became 70.

For C1 and C2 levels, the entire dataset was used and the number of distinct classes was 2 and 6, respectively.

## 5 Results

This section includes an overview of the results achieved by all of the methods presented in Section 3. Before delving into the detailed performance for each method, in Table 2 we report the running time and memory footprint needed to evaluate the embedding on the full dataset. Experiments do not include the evaluation of the distance matrix and/or the clustering algorithm solution.

**Table 2:** Efficiency of the 14 strategies to calculate the embedding/kernel matrix. Results have obtained on a 14” MacBook Pro with 16-cores M4 Max CPU and 128GB of RAM.

| Family | Method | Running Time | Memory [MB]<br>avg (peak) |
| --- | --- | --- | --- |
| BoW-inspired | BoH | 3 [s] | 1,169 (1,857) |
|  | BoN w/ Betweenness Centrality<sup>†</sup> | 395 [s] | 22,413 (22,969) |
|  | BoN w/ Degree Centrality<sup>†</sup> | 5 [s] | 10,310 (22,992) |
|  | BoN w/ Node Degree | 3 [s] | 1,365 (1,892) |
|  | BoN w/ PageRank<sup>†</sup> | 6 [s] | 11,876 (23,080) |
| Kernel-based | Histogram Kernel | 2 [s] | 909 (1,512) |
|  | Jaccard Kernel<sup>†</sup> | 34 [s] | 12,042 (17,491) |
|  | Edit Kernel<sup>†</sup> | 29 [h] | 20,500 (20,785) |
|  | Stratified Edit Kernel<sup>†</sup> | 30 [h] | 20,605 (20,892) |
| Neural Networks-based | Graph2Vec<sup>‡</sup> | 18 [s] | 6,301 (10,131) |
|  | HGAE (Complete)<sup>‡</sup> | 610 [s] | 19,699 (19,981) |
|  | HGAE (Hyperedges-Only)<sup>‡</sup> | 463 [s] | 19,588 (19,907) |
|  | HGAE (One-Hot+Hyperedges)<sup>‡</sup> | 995 [s] | 20,005 (20,176) |

<sup>†</sup>These methods exploit multiprocessing strategies to parallelize execution. In our experiments, the number of processes equals the number of CPU cores.
<sup>‡</sup>Running times and memory occupation refer to a fully CPU-based implementation (no GPU acceleration) and they include both training and inference.

### 5.1 BoN and BoH Methods Results

Figure 1 illustrates the performances of the BoW-inspired methods across all 4 classification levels (panels 1a–1d, respectively) as a function of the number of clusters (i.e., the cut-point on the hierarchical clustering dendrogram). The vertical dashed line in each subplot represents the number of distinct classes present in each classification level (in our dataset). From the plots it is apparent how methods using hyperedges as pivotal features for the embedding of the hypergraphs tend to have higher performances compared to BoN methods on all classification levels. This result appears as a consequence of the fact that the hyperedge set of a metabolic hypergraph exactly coincides with the set of chemical relations happening inside an organism, the latter being a fingerprint of its taxonomy. The overall best performer seems to be the BoH embedding with Jaccard distance reaching homogeneity scores *h* ≥ 0.8 for all classification levels. In particular the score for C1 was slightly above 0.95, highlighting an almost perfect clustering discrimination.

**Fig. 1:**
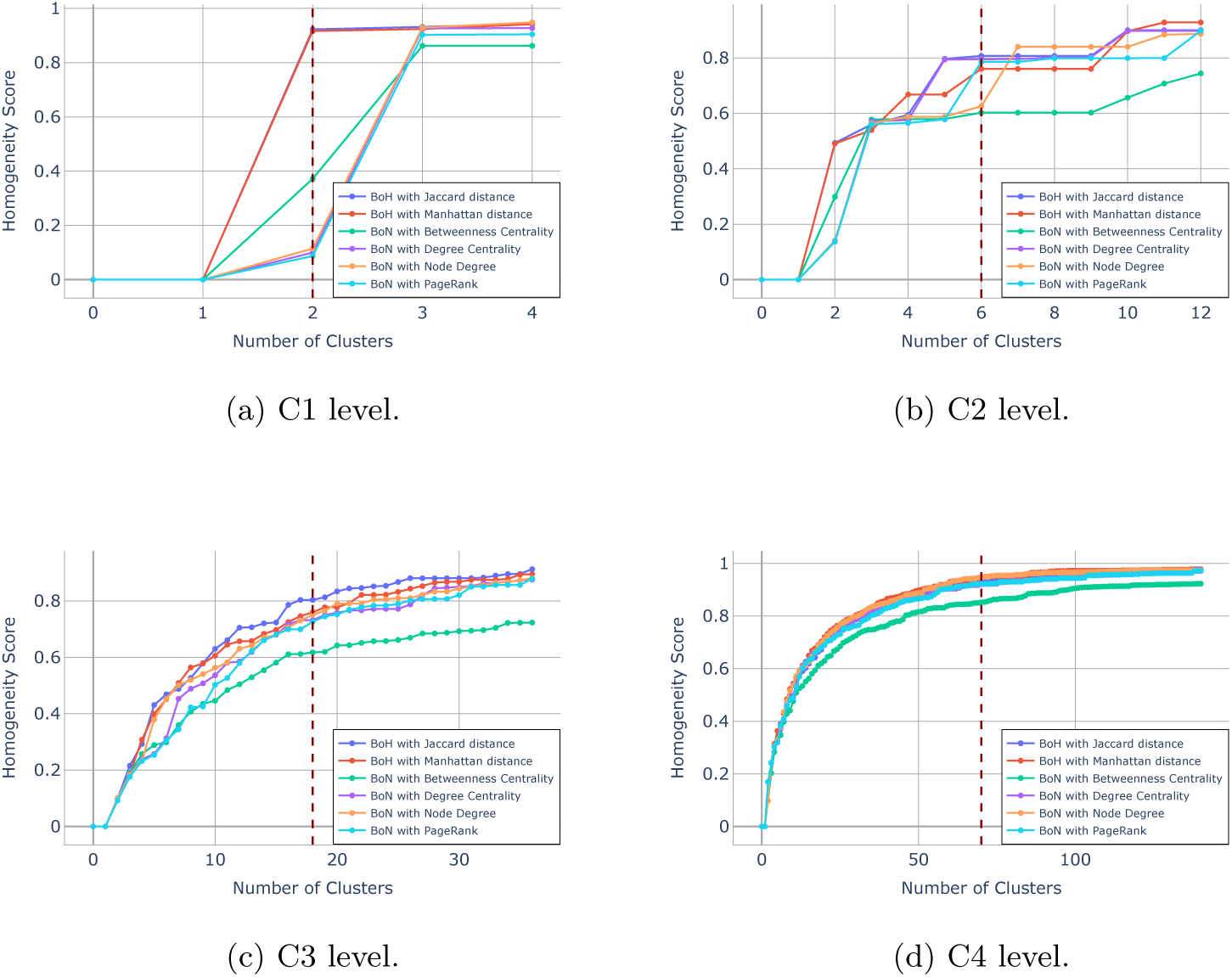
Homogeneity score vs. number of clusters for BoW inspired methods.

### 5.2 Kernel Methods Results

Figure 2 illustrates the performances of the kernel methods on the 4 classification levels (panels 2a–2d, respectively). The HCK and the JK perform significantly better than the ones based on Edit distances. In particular, HCK manages to reach homogeneity *h* = 1 on the first classification level keeping the number of clusters equal to the number of classes, meaning that Eukaryotes and Prokaryotes are perfectly distinguished using the HCK. The SEK in particular shows lack of performances probably due to the abundance of hyperedges of order two in the metabolic hypergraphs. Giving the same relevance to all hyperedge orders (i.e. sizes) was probably not a suitable approach for this particular application. The performances overall are a bit superior to the ones obtained by BoW inspired methods.

**Fig. 2:**
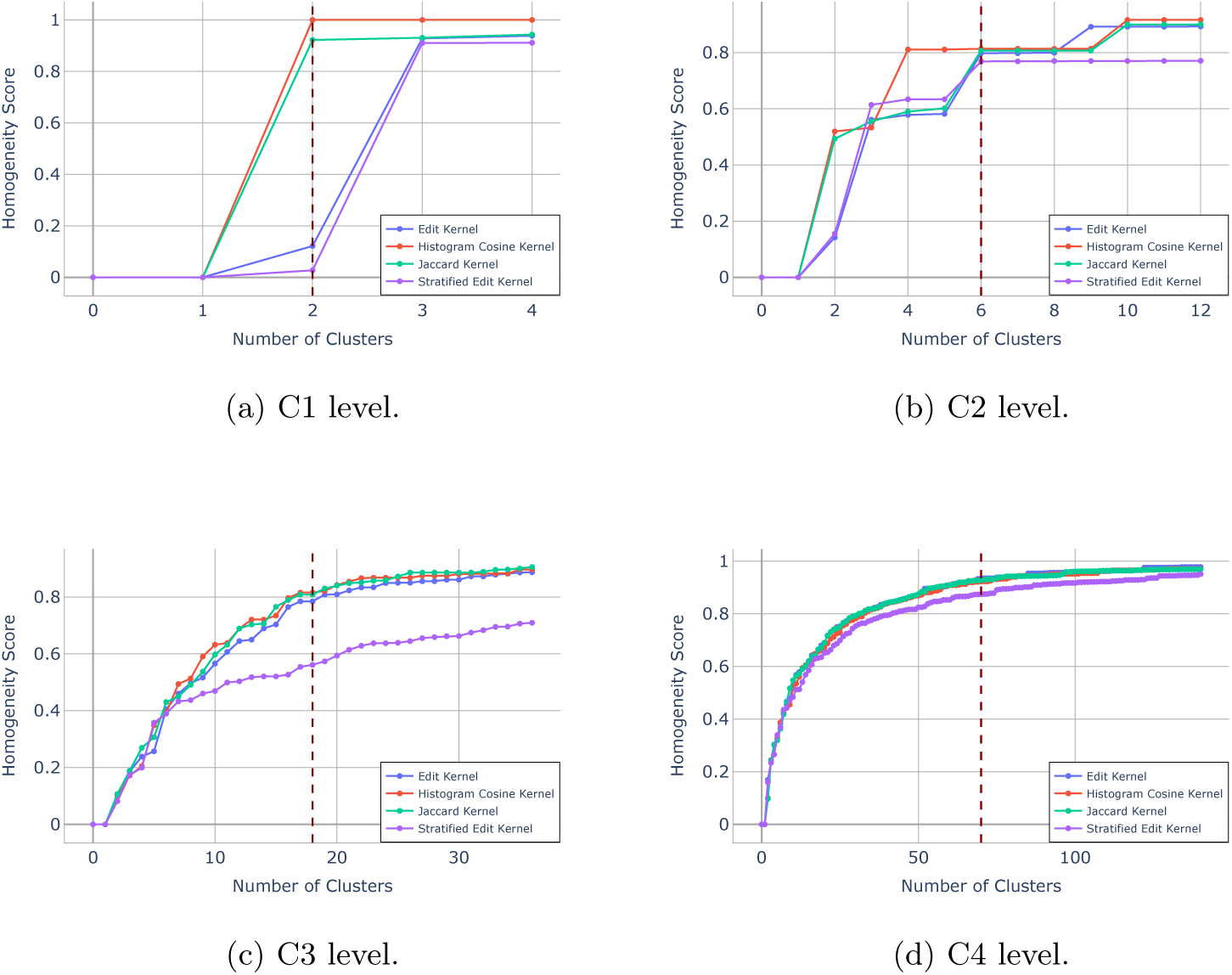
Homogeneity score vs. number of clusters for kernel methods.

### 5.3 Neural Network-Based Methods Results

Figure 3 illustrates the performances of the neural network-based methods on the 4 classification levels (panels 3a–3d, respectively). Graph2Vec and the HGAE-Hyperedges-Only display the same perfect performance as the HCK on C1 which is very promising. Considering the other classification levels, the HGAE-Hyperedges-Only exhibits good performances in all of them while the Graph2Vec shows a slightly lower performance overall. Both HGAE-Complete and HGAE-One-Hot+Hyperedges structures perform generally worse than the one considering only hyperedges. This phenomenon suggests that information regarding the chemical reactions of an organism (i.e., hyperedges) is the most relevant for the creation of effective hypergraph level embeddings for metabolic networks. Convolution also does not seem to be optimal probably due to metabolic networks being extremely sparse hypergraphs.

**Fig. 3:**
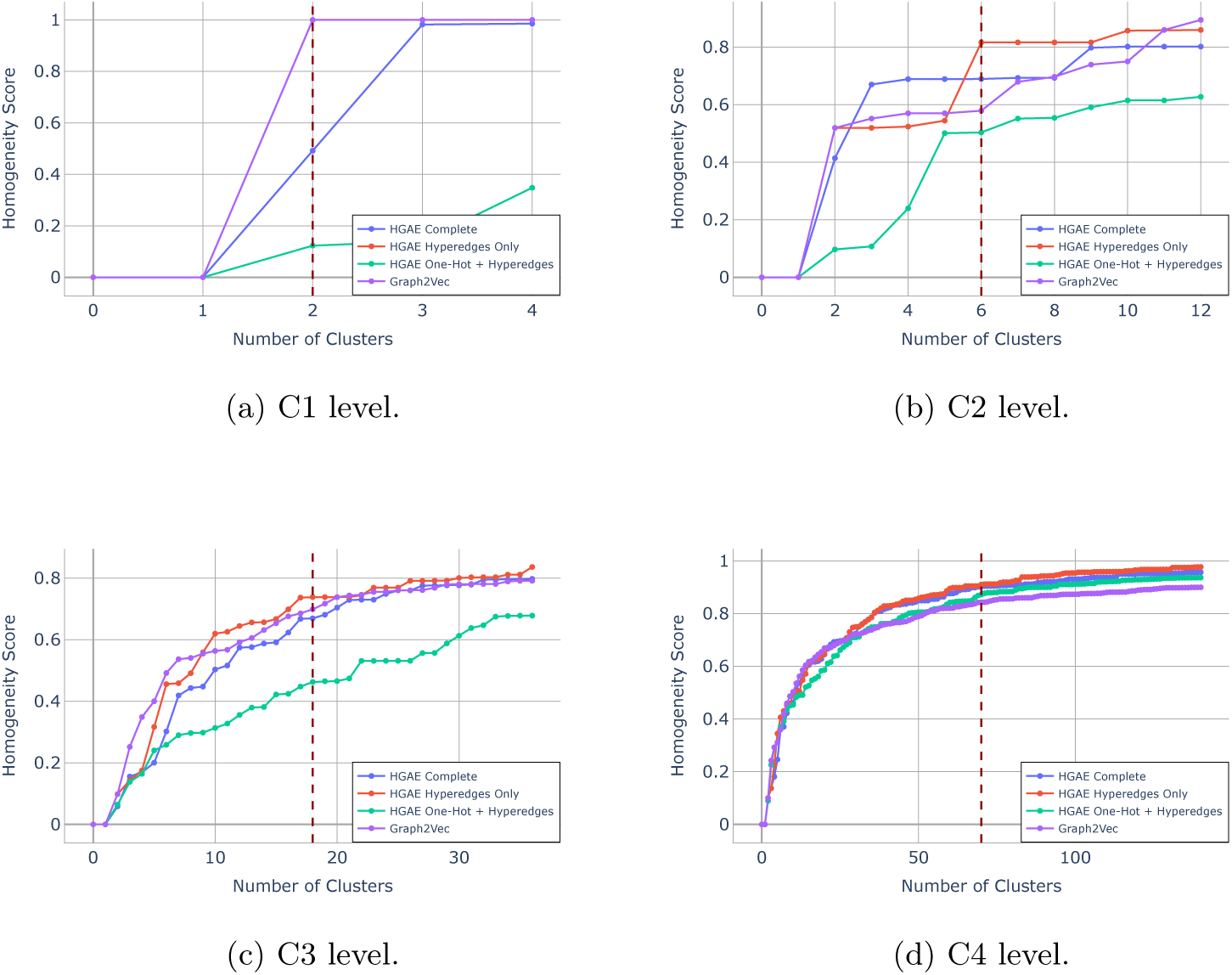
Homogeneity score vs. number of clusters for neural network-based methods.

### 5.4 Best Performers Results

This section provides the detailed analysis of the clustering solutions deriving from the best performing method from each of the families analyzed in the previous sections. The best performers were chosen according to their performance in terms of overall homogeneity score (cfr. Figures 1–3), yielding the following shortlist: BoH with Jaccard distance (for BoW-inspired methods), HCK (for kernel-based methods) and HGAE-Hyperedges-Only (for neural network-based methods). A first important observation is that all of the best performing models create the embeddings starting directly from the hypergraphs of the metabolic networks. This finding suggests that embeddings deriving from clique expansions tend to loose important details and therefore are less informative compared to methodologies which are natively tailored to the hypergraph nature of metabolic networks. Conversely to the previous results from Sections 5.1–5.3 –in which we have shown the homogeneity scores as a function of the number of clusters– we are now interested in analyzing, for each best performer, the resulting best clustering solution in terms of clusters composition. As anticipated in Section 4, in order to determine the appropriate number of clusters for each classification level and each embedding method, the silhouette score was used. Specifically, the chosen solution was the one that maximized the silhouette score in a symmetric neighborhood of the number of distinct classes at each classification level.

Figure 4 presents the results on level C1 and, in accordance with the very high homogeneity scores presented in the previous sections, both the HCK and HGAE-Hyperedges-Only manage to perfectly separate eukaryotic and prokaryotic organisms (i.e., two clusters with maximal purity). Also the BoH approach manages to achieve almost perfect separation with a minimum purity score among the clusters of 0.996. In conclusion, the selected embedding methodologies are explicative enough to represent the first level of the Linnaeus taxonomy perfectly.

**Fig. 4:**
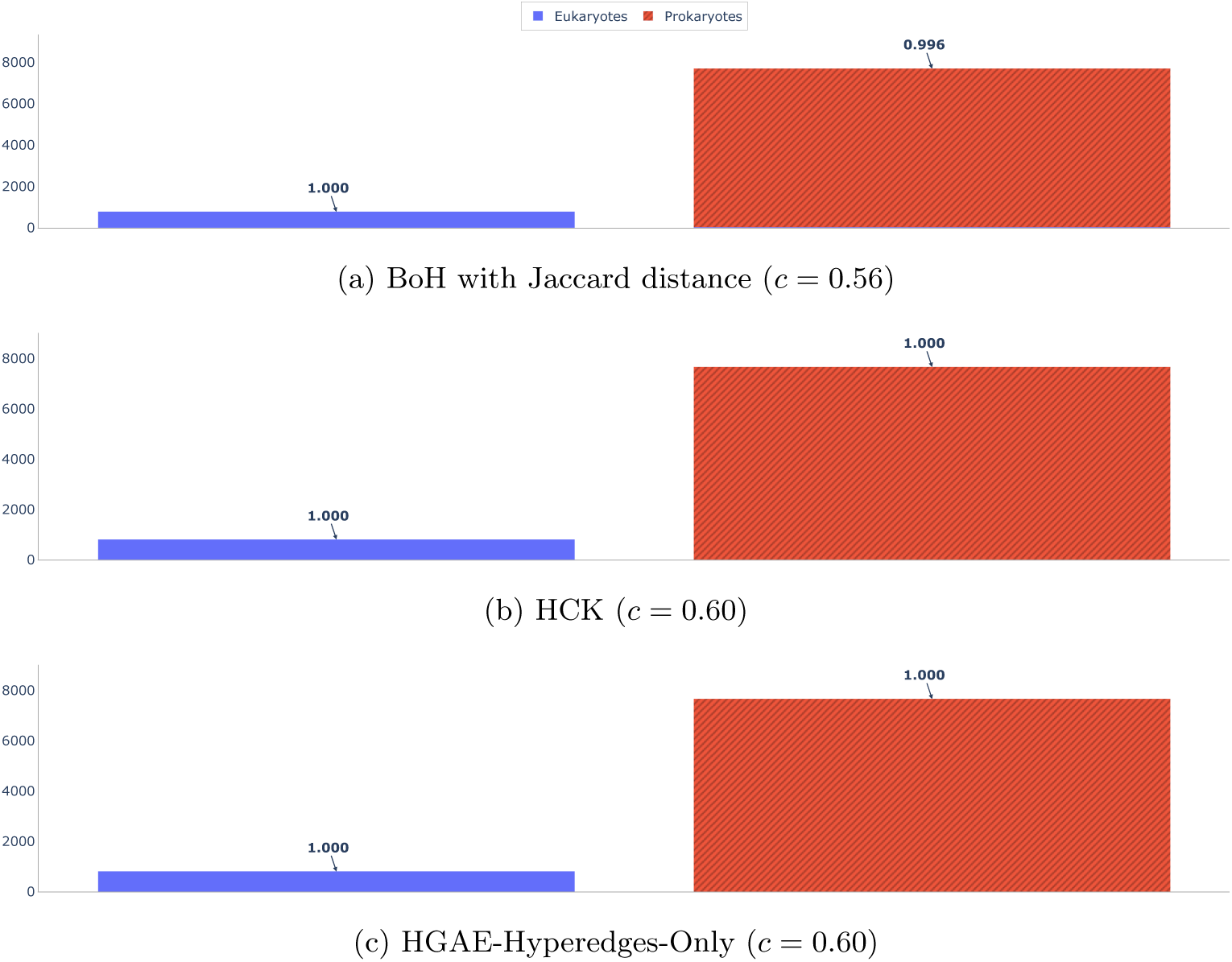
Clustering solutions of the best performers for level C1. For each clustering solution we show as many bars as there are clusters, with the height of each bar showing the number of samples in each cluster. Bars are further colored according to the ground-truth class (see legend) and for each cluster we also report the purity score *p*. Finally, for each clustering solution, we also show the cophenetic correlation *c*.

Figure 5 presents the results on C2 level, in this case different embedding methodologies maximized their silhouette score at a different number of clusters inside the candidate range, which for C2 was [4, 8]. With a total of 6 distinct classes in C2, the HCK and the HGAE-Hyperedges-Only had the most conservative result, with only 4 clusters while the BoH highlighted 6 clusters. Despite the differences in the number of clusters all solutions tend to present the same limitation: Bacteria tend to split up in different (albeit pure, *p >* 0.8 for all methods) clusters while the Eukaryotic organisms (i.e., Animals, Fungi, Plants and Protists) remain in a single, separate, and less pure cluster (*p* ≈ 0.6 for all methods). The HGAE was not able to accurately isolate Archea however it managed to keep most of the Bacteria into a single cluster. This phenomenon highlights how, in all the presented embedding spaces, different kinds of Bacteria are more widely separated from one another than different kingdoms of Eukaryotic organisms are from each other. This result is probably due to Bacteria representing the majority of the dataset and presenting many differences among them, also considering their effect on Humans: from rare and very dangerous (e.g., *Bacillus Anthracis*) to common and harmless (e.g., *Staphylococcus Epidermidis*).

**Fig. 5:**
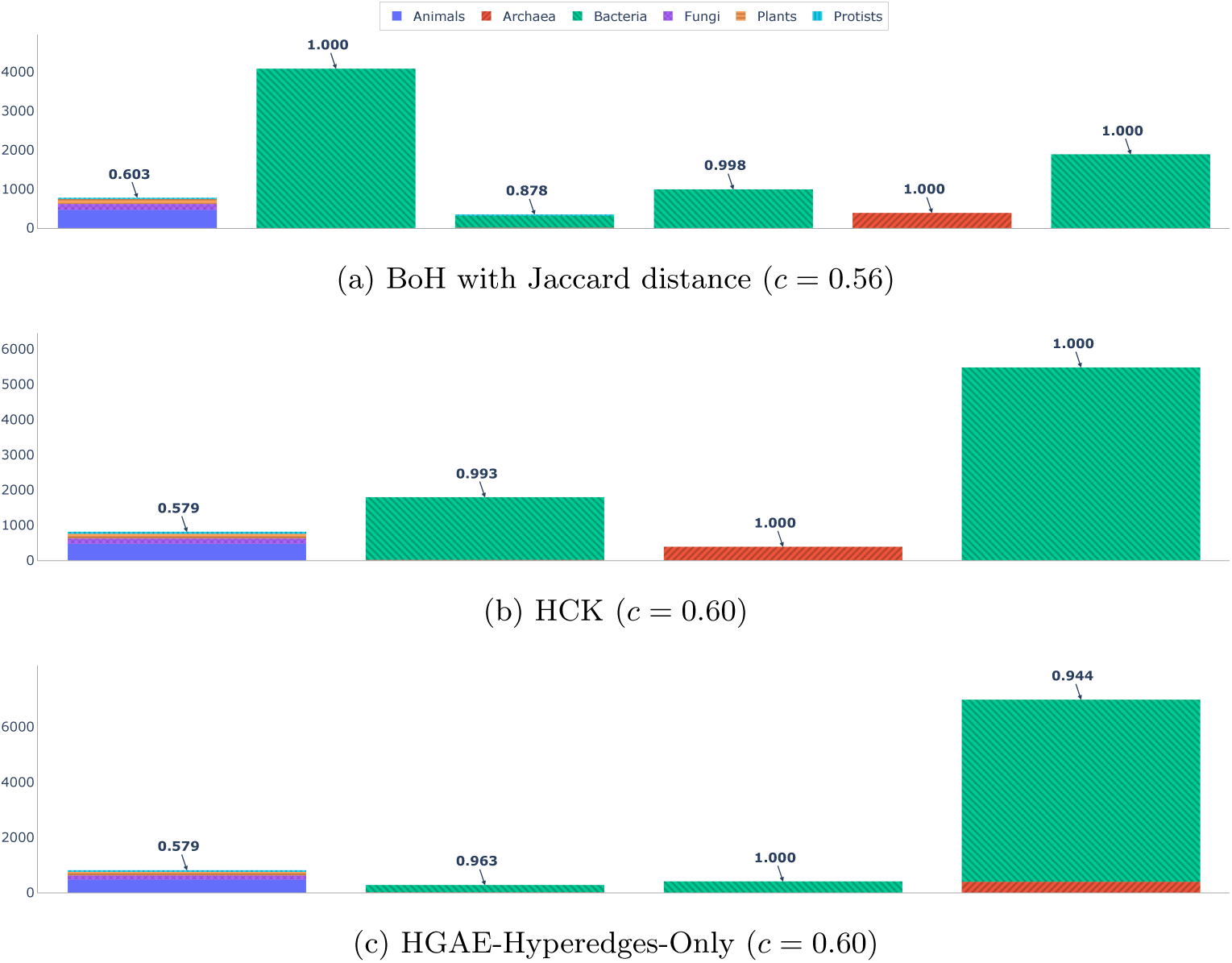
Clustering solutions of the best performers for level C2. For details, see caption of Fig. 4.

When considering C3 level, presented in Figure 6, the HGAE-Hyperedges-Only highlighted 21 clusters while the other methods settled on 23 clusters, the allowed range for C3 was [13, 23]. As anticipated in Section 5 the number of distinct classes in C3 was lowered to 18 by cutting any class that did not include at least 100 organisms in the dataset. Results were overall coherent among all clusters: the same classes tend to split up in different clusters for all embedding spaces (e.g, *Firmicutes - Bacilli* and *Actinobacteria*). All methods exhibit a very similar performance however the separation is far from perfect, all methods show between 15 and 18 clusters with very high purity while the rest remains quite mixed with scores ranging from *p* = 0.4 to *p* = 0.7. The HGAE in particular includes 7 very small clusters in its clustering solution while other methods show only 2 or 3 micro clusters.

**Fig. 6:**
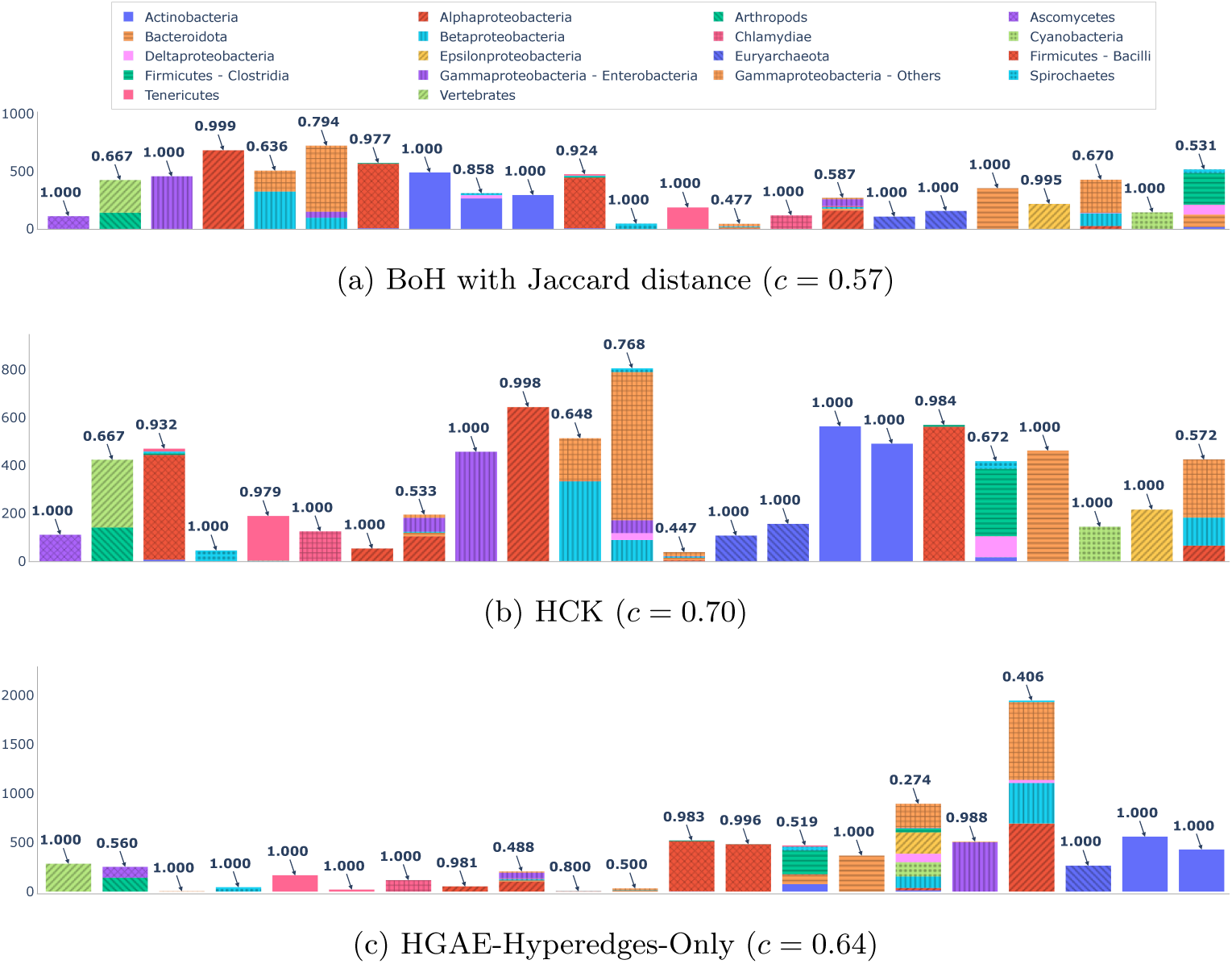
Clustering solutions of the best performers for level C3. For details, see caption of Fig. 4.

For C4 level, presented in Figure 7, the number of clusters differs again just as in the case of C3 level, the search range in this case was [40, 100] considering a total of 70 remaining distinct class labels after cutting any class not including at least 20 organisms. In this case the HGAE-Hyperedges-Only and BoH selected the lowest number of clusters (82) while the HCK went for 85. The type of visualization adopted in Figure 7 for C4 level is different compared to the other classification levels, in order to accommodate for the high number of classes and clusters. Indeed, each bar shows in green the majority class in a certain cluster and in red the sum of all minority classes in the same cluster. The numeric purity score was removed as well to foster readability. The performances on C4 level are exceptionally remarkable, with the vast majority of the clusters exhibiting a very high degree of purity for all embedding methods. While BoH and the HCK show only one cluster with major impurity, the HGAE highlights more clusters with relatively low purity. This is probably due to the fact that HGAE embeddings derive directly from Neural Layers which (as explained in Section 3.4) are intended to minimize reconstruction loss without explicitly defining which aspects of the structures drive latent representations.

**Fig. 7:**
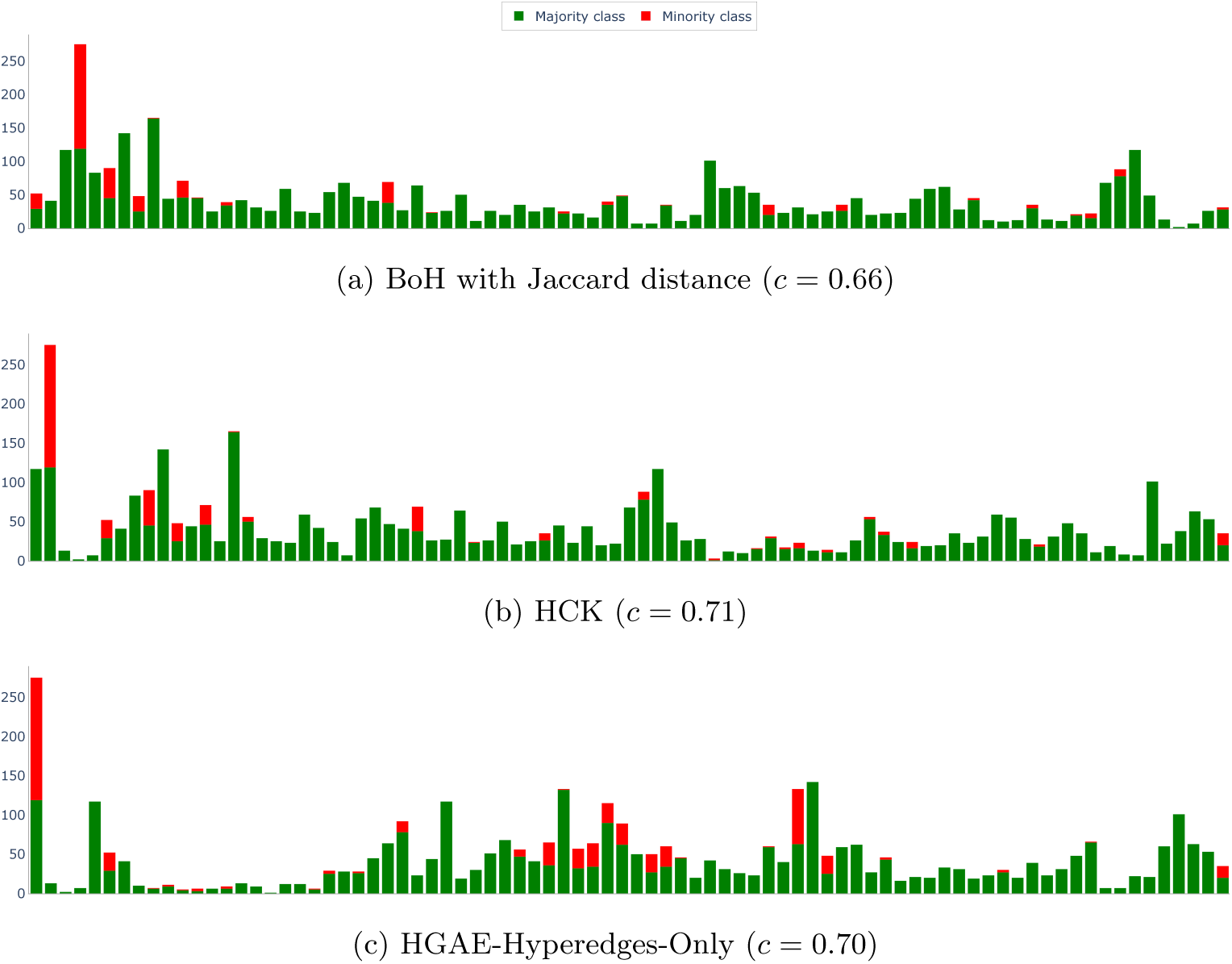
Clustering solutions of the best performers for level C4. For details, see caption of Fig. 4, with the only modification that –due to the high number of classes– we show, for each cluster, the composition in terms of number of organisms in the majority class vs. minority class (see legend).

In Figures 8–10, we show the UMAP visualizations [60, 61] of the three best performing embedding strategies for each of the classification levels from C1 to C3 (C4 was excluded due to the high number of distinct classes which made the plots unreadable). Figure 8 highlights a very clear separation of the classes in the UMAP dimensions at the C1 level. At the C2 level (Figure 9), the separation is less precise however definitely present. Specifically most classes appear in very concentrated hot-spots, Eucaryotic organisms however tend to mix together. This result strengthens the previous observation on the diversity of Bacteria compared to other C2 classes. When looking at C3 (Figure 10), the results are mitigated as some classes are extremely well isolated (e.g., *Alphaproteobacteria*) while others appear quite mixed (e.g., *Gammapro-teobacteria - Enterobacteria*, *Betaproteobacteria*, *Firmicutes - Clostridia*, etc.). It is important to note that these figures include only organisms that were considered during the clustering of C3. These visualizations overall confirm the results presented in this section.

**Fig. 8:**
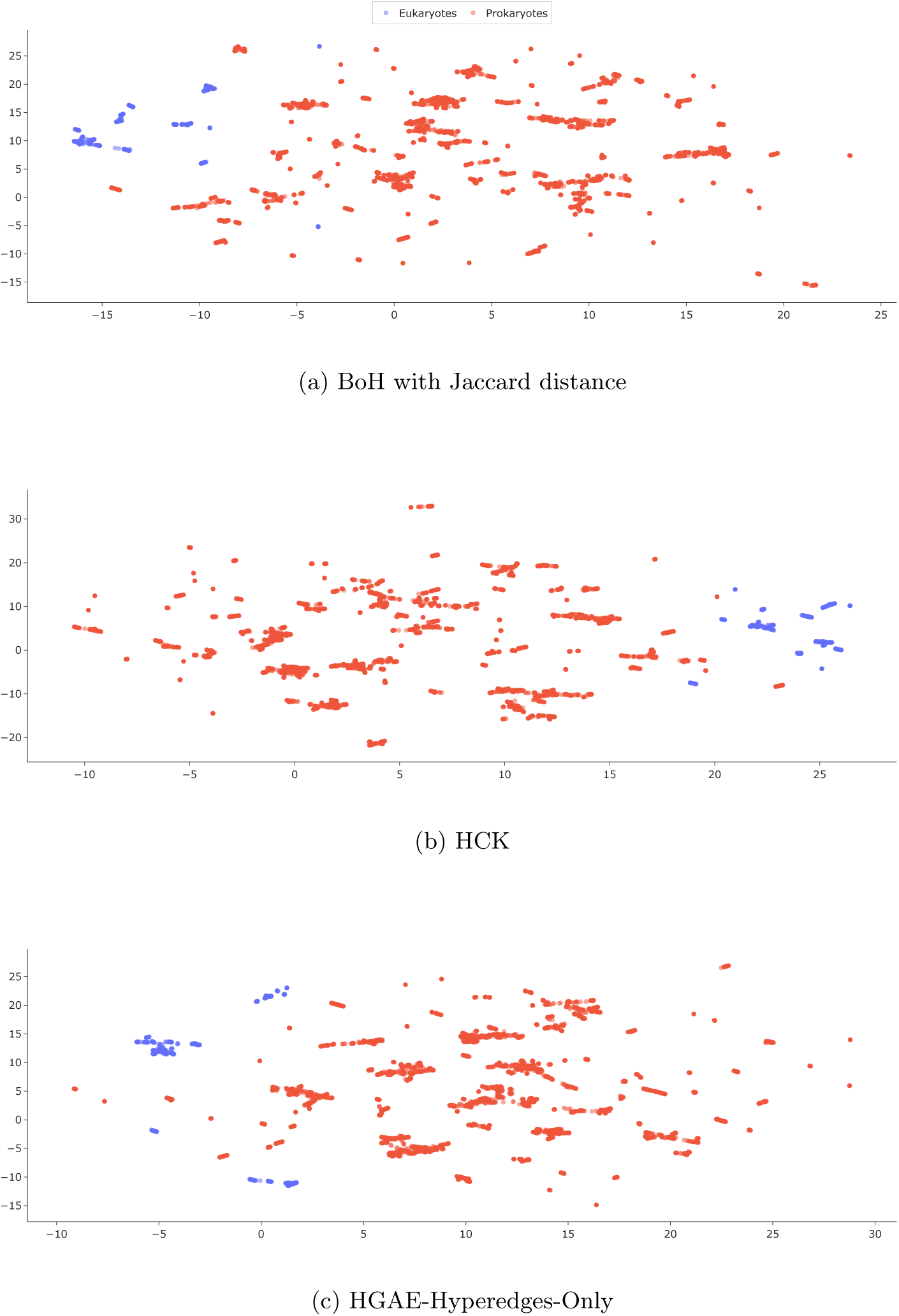
2D UMAP visualization of the best performers for level C1.

**Fig. 9:**
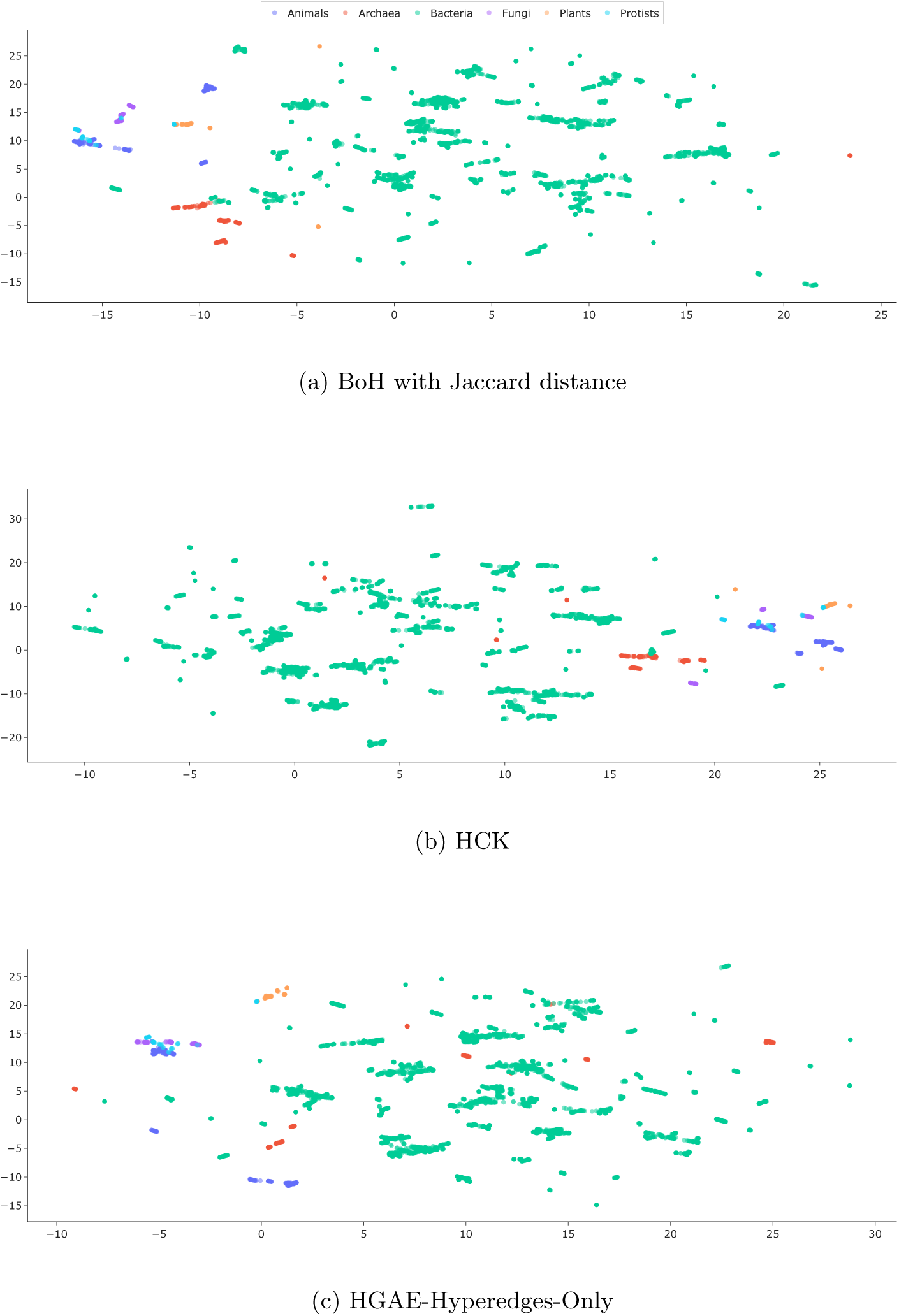
2D UMAP visualization of the best performers for level C2.

**Fig. 10:**
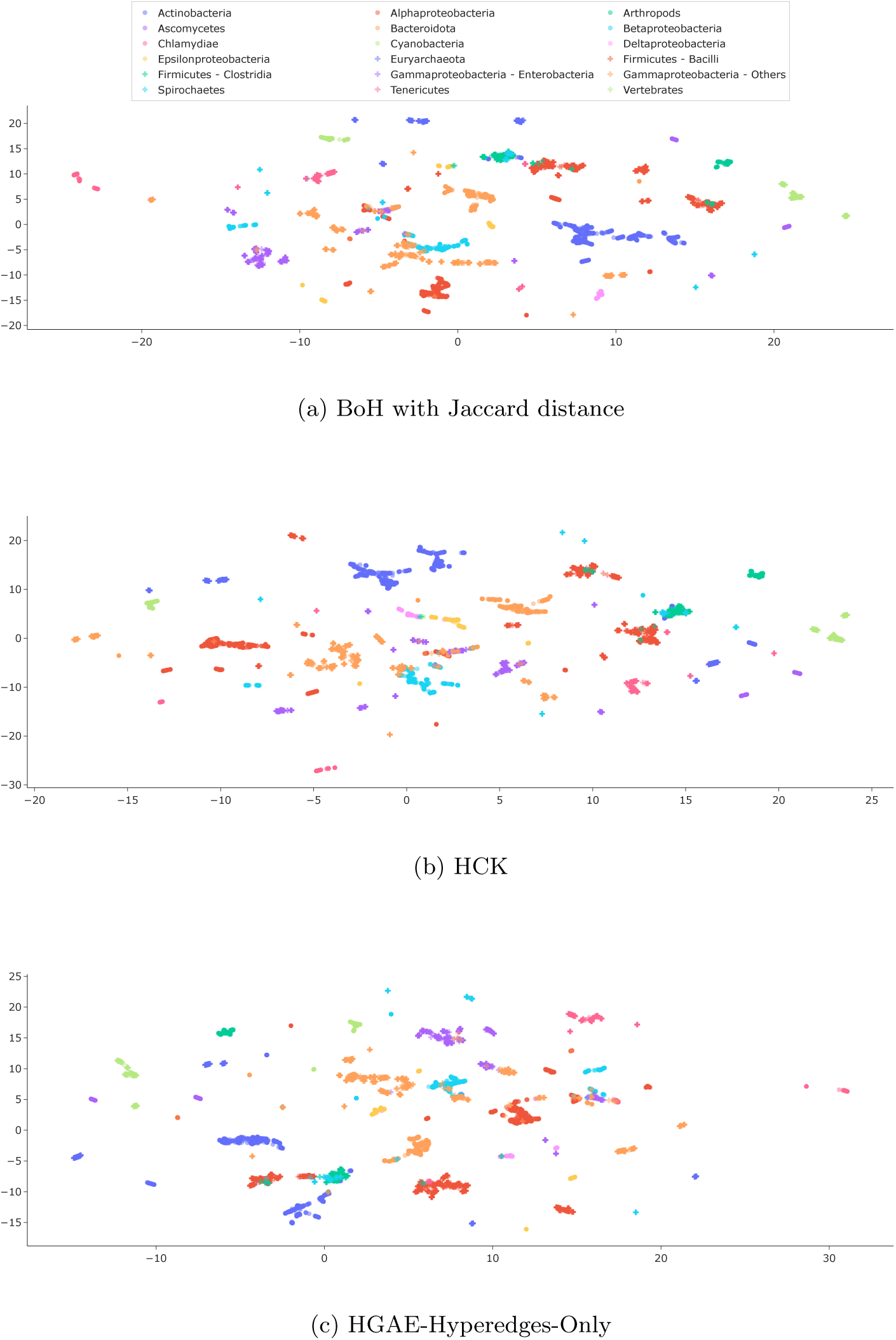
2D UMAP visualization of the best performers for level C3.

Finally, to complete the overview of the separation between organisms, Figures 11–13 show the heatmaps of the distance matrices calculated from the three best performing embedding methods. For each best performer, panels (a), (b), (c) and (d) refer to C1, C2, C3 and C4, respectively, with red solid lines grouping together organisms pertaining to different classes at each taxonomic level.

**Fig. 11:**
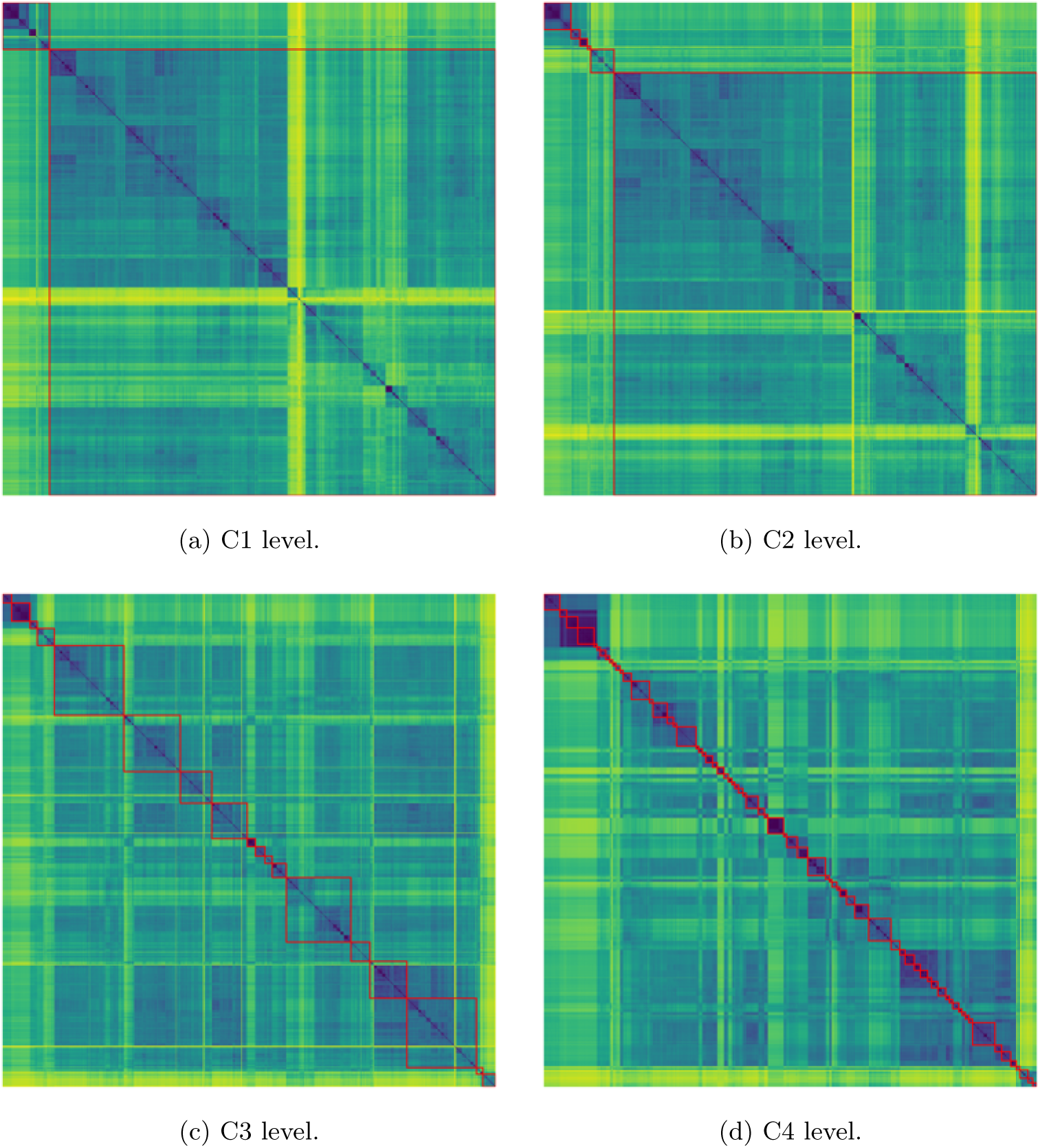
Distance heatmap (the lighter, the farther) obtained by BoH with Jaccard distance on the four taxonomic levels.

**Fig. 12:**
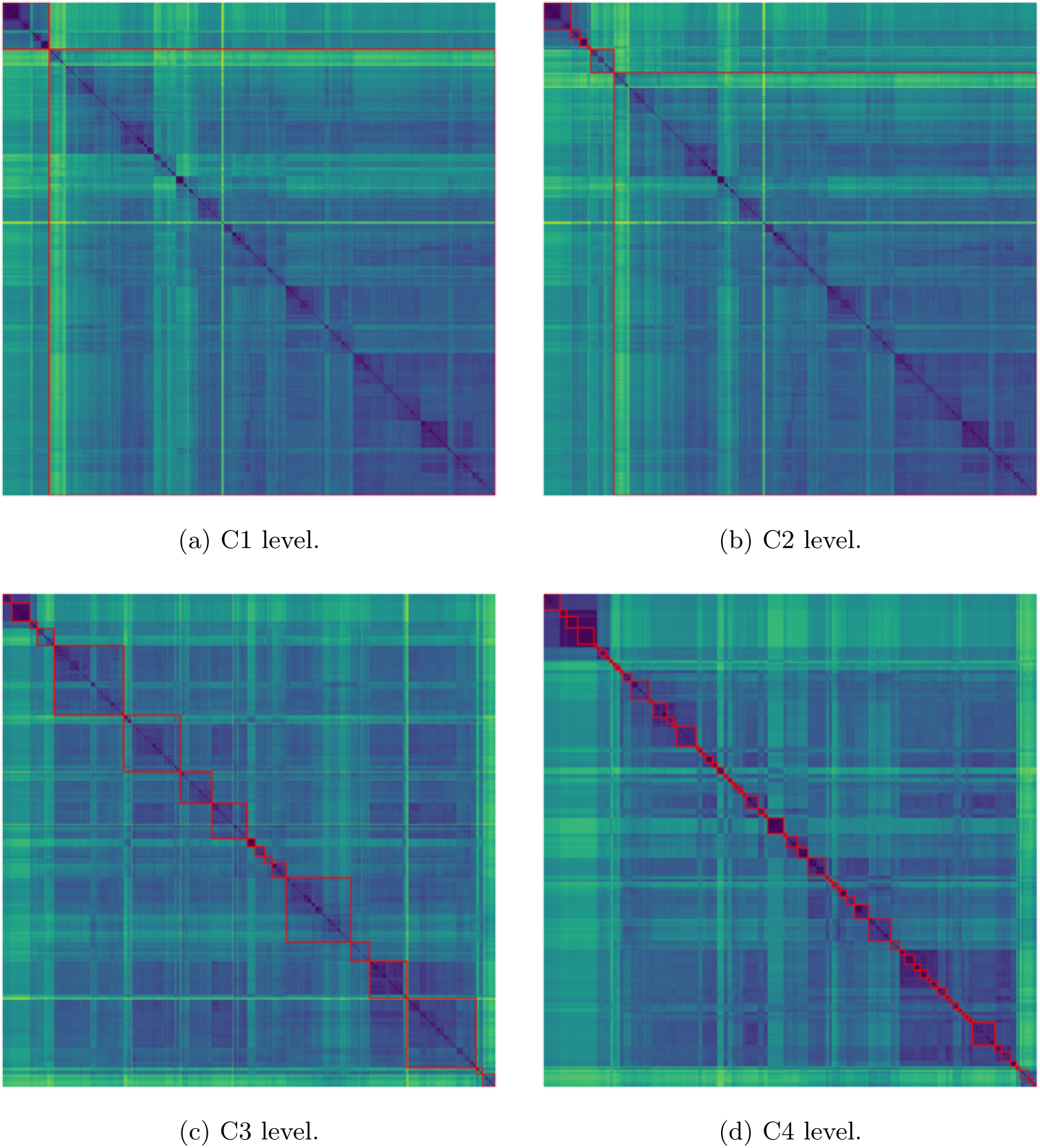
Distance heatmap (the lighter, the farther) obtained by HCK on the four taxonomic levels.

**Fig. 13:**
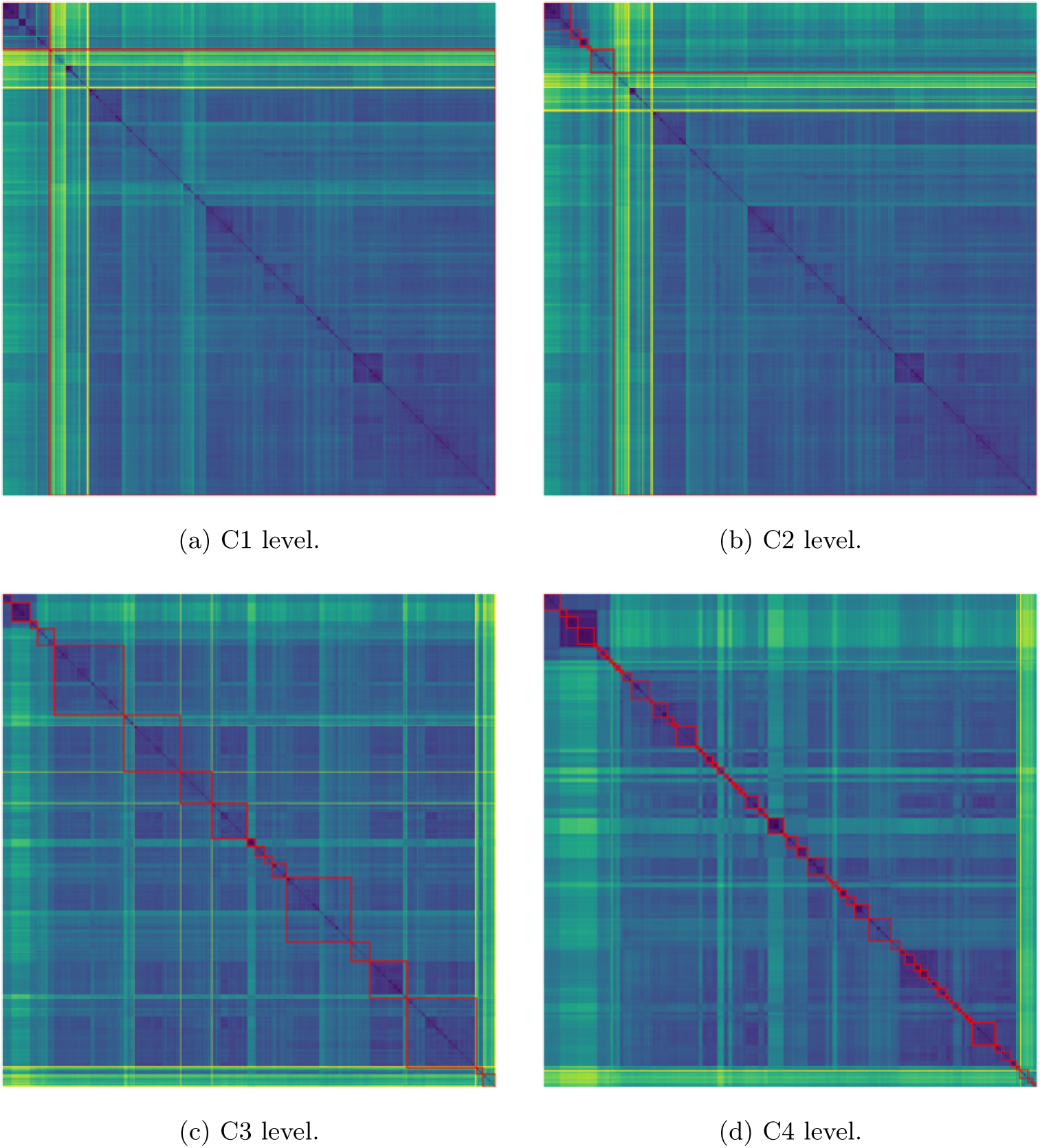
Distance heatmap (the lighter, the farther) obtained by HGAE-Hyperedges-Only on the four taxonomic levels.

## 6 Conclusions and perspectives

In this paper, we have investigated the problem of embedding in a vectorial space sets of metabolic reactions, originally represented as hypergraphs of compounds, in order to perform unsupervised clustering of organisms. We have presented a wide range of embedding methodologies, spanning from Bag-of-Words inspired methods over hyper-edges and nodes (easy to interpret and to compute, but potentially sparse and high dimensional), to Kernel-based methods (which implicitly embed the data points and are not interpretable) and Neural Network-based methods (which are compact and dense, but not interpretable).

The embedding methodologies were then used to perform unsupervised clustering of organisms at different levels of Linnaeus’ taxonomy, from the most coarse-grained (C1 level, that is, cellular organization) to the most fine-grained (C4 level, that is, genus), and they have been evaluated in terms of homogeneity (at clustering solution level) and purity (at individual cluster level) of the resulting clusters.

The overall performances of the methods remain quite high across all levels of Linnaeus’ taxonomy, with C3 and C2 levels presenting the most challenges. Future embedding methodologies should probably be developed especially to face the intricacies of those classification levels. Besides that, considering the fully unsupervised clustering framework, the results are satisfactory and open the possibility for additional research on the optimal embedding space for metabolic hypergraphs.

Finally, we advocate that the embedding method should be chosen based on the goal of the downstream task to be performed and believe our findings will be of interest to researchers working on computational biology, metabolic modeling, and network-based analysis of evolutionary systems. Beyond taxonomy recovery, such embeddings could support tasks like metabolic pathway reconstruction, prediction of missing reactions, identification of functionally similar organisms, or analysis of host–microbiome metabolic complementarity. In these contexts, it would be important not only to develop task-specific evaluation criteria, but also to design embedding strategies tailored to the biological structure and functional goals of each task.

We mention that we considered undirected graphs and hypergraphs, thus losing important information about reaction paths and causality. Future work could explore the possibility to consider embedding methods for directed interactions, and / or taking into account stoichiometry coefficients (for e.g. authorizing multiset hyperedges in the context of hypergraphs).

## Declarations

## Ethics approval and consent to participate

Not applicable.

## Consent for publication

Not applicable.

## Data availability

The metabolic data supporting the conclusions of this article are available in the KEGG repository at https://www.kegg.jp. The list of 8,467 organisms is included within the article as a supplementary material.

## Materials availability

Not applicable.

## Code availability

The code reproducing all embedding routines is available at https://github.com/Alessi0X/HypergraphEmbedding4MetabolicNetworks.

## Author contribution

A.M., B.S. and C.M. conceived the idea and the research questions. A.M. took care of data collection, selected the methods to be tested and supervised the work. M.C. implemented all tested methods, conceiving the proposed ones, and performed the data analysis. M.C., B.S., C.M. and A.M. wrote the main manuscript. All authors reviewed the manuscript and agreed with its submission.

## Footnotes

1 https://www.kegg.jp/brite/br08601

## References

[1] Jeong, H., Tombor, B., Albert, R., Oltvai, Z.N., Barabási, A.-L.: The large-scale organization of metabolic networks. Nature 407(6804), 651–654 (2000) 10.1038/35036627

[2] Galhardo, M., Sinkkonen, L., Berninger, P., Lin, J., Sauter, T., Heinäniemi, M.: Integrated analysis of transcript-level regulation of metabolism reveals disease-relevant nodes of the human metabolic network. Nucleic Acids Research 42(3), 1474–1496 (2013) 10.1093/nar/gkt989

[3] O’Brien, E., Monk, J., Palsson, B.: Using genome-scale models to predict biological capabilities. Cell 161(5), 971–987 (2015) 10.1016/j.cell.2015.

[4] Klamt, S., Haus, U.-U., Theis, F.: Hypergraphs and cellular networks. PLOS Computational Biology 5(5), 1–6 (2009) 10.1371/journal.pcbi.1000385

[5] Ritz, A., Tegge, A.N., Kim, H., Poirel, C.L., Murali, T.M.: Signaling hypergraphs. Trends in Biotechnology 32(7), 356–362 (2014) 10.1016/j.tibtech.2014.04.007

[6] Estrada, E., Rodríguez-Velázquez, J.A.: Subgraph centrality and clustering in complex hyper-networks. Physica A: Statistical Mechanics and its Applications 364, 581–594 (2006) 10.1016/j.physa.2005.12.002

[7] Arodź, T.: Clustering organisms using metabolic networks. In: Proceedings of the 8th International Conference on Computational Science, Part II. ICCS ’08, pp. 527–534. Springer, Berlin, Heidelberg (2008). 10.1007/978-3-540-69387-1_60

[8] García, I., Chouaia, B., Llabrés, M., Simeoni, M.: Exploring the expressiveness of abstract metabolic networks. PLOS ONE 18(2), 1–27 (2023) 10.1371/journal.pone.0281047

[9] Kirkland, S.: Two-mode networks exhibiting data loss. J. Complex Netw. 6(2), 297–316 (2018) 10.1093/comnet/cnx039

[10] Zien, J.Y., Schlag, M.D.F., Chan, P.K.: Multilevel spectral hypergraph partitioning with arbitrary vertex sizes. IEEE Transactions on Computer-Aided Design of Integrated Circuits and Systems 18(9), 1389–1399 (1999) 10.1109/43.784130

[11] Giamphy, E., Guillaume, J.-L., Doucet, A., Sanchis, K.: A survey on bipartite graphs embedding. Social Network Analysis and Mining 13(1), 54 (2023) 10.1007/s13278-023-01058-z

[12] Maleki, S., Saless, D., Wall, D.P., Pingali, K.: HyperNetVec: Fast and Scalable Hierarchical Embedding for Hypergraphs, pp. 169–183. Springer, Cham (2022). 10.1007/978-3-030-97240-0_13

[13] Cocco, N., Llabrés, M., Reyes-Prieto, M., Simeoni, M.: MetNet: A two-level approach to reconstructing and comparing metabolic networks. PLOS ONE 16(2), 1–18 (2021) 10.1371/journal.pone.0246962

[14] Zhu, D., Qin, Z.S.: Structural comparison of metabolic networks in selected single cell organisms. BMC Bioinformatics 6(1), 8 (2005) 10.1186/1471-2105-6-8

[15] Palmer-Rodríguez, P., Alberich, R., Reyes-Prieto, M., Castro, J.A., Llabrés, M.: Metadag: a web tool to generate and analyse metabolic networks. BMC Bioinformatics 26(1), 31 (2025) 10.1186/s12859-025-06048-w

[16] Tun, K., Dhar, P.K., Palumbo, M.C., Giuliani, A.: Metabolic pathways variability and sequence/networks comparisons. BMC Bioinformatics 7(1), 24 (2006) 10.1186/1471-2105-7-24

[17] Ebenhöh, O., Handorf, T.: Functional classification of genome-scale metabolic networks. EURASIP Journal on Bioinformatics and Systems Biology 2009, 1–13 (2009) 10.1155/2009/570456

[18] Reyes, J., Dunkel, J.: Functional classification of metabolic networks. arXiv preprint (2025) 2503.14437

[19] Ramon, C., Stelling, J.: Functional comparison of metabolic networks across species. Nature Communications 14(1), 1699 (2023) 10.1038/s41467-023-37429-5

[20] Martino, A., Giuliani, A., Todde, V., Bizzarri, M., Rizzi, A.: Metabolic networks classification and knowledge discovery by information granulation. Computational Biology and Chemistry 84, 107187 (2020) 10.1016/j.compbiolchem.2019.107187

[21] Granata, I., Guarracino, M.R., Kalyagin, V.A., Maddalena, L., Manipur, I., Pardalos, P.M.: Supervised classification of metabolic networks. In: 2018 IEEE International Conference on Bioinformatics and Biomedicine (BIBM), pp. 2688– 2693 (2018). 10.1109/BIBM.2018.8621500

[22] Granata, I., Guarracino, M.R., Kalyagin, V.A., Maddalena, L., Manipur, I., Pardalos, P.M.: Model simplification for supervised classification of metabolic networks. Annals of Mathematics and Artificial Intelligence 88(1), 91–104 (2020) 10.1007/s10472-019-09640-y

[23] Mueller, L.A., Kugler, K.G., Netzer, M., Graber, A., Dehmer, M.: A network-based approach to classify the three domains of life. Biology Direct 6(1), 53 (2011) 10.1186/1745-6150-6-53

[24] Kanehisa, M., Goto, S.: KEGG: Kyoto encyclopedia of genes and genomes. Nucleic acids research 28(1), 27–30 (2000) 10.1093/nar/28.1.27

[25] Cokelaer, T., Pultz, D., Harder, L.M., Serra-Musach, J., Saez-Rodriguez, J.: BioServices: a common Python package to access biological Web Services programmatically. Bioinformatics 29(24), 3241–3242 (2013) 10.1093/bioinformatics/btt547

[26] Cock, P.J.A., Antao, T., Chang, J.T., Chapman, B.A., Cox, C.J., Dalke, A., Friedberg, I., Hamelryck, T., Kauff, F., Wilczynski, B., Hoon, M.J.L.: Biopython: freely available Python tools for computational molecular biology and bioinformatics. Bioinformatics 25(11), 1422–1423 (2009) 10.1093/bioinformatics/btp163

[27] Federhen, S.: The NCBI Taxonomy database. Nucleic Acids Research 40(D1), 136–143 (2011) 10.1093/nar/gkr1178

[28] Martino, A., Rizzi, A.: (Hyper)graph Kernels over Simplicial Complexes. Entropy 22(10), 1155 (2020) 10.3390/e22101155

[29] Ward Jr., J.H.: Hierarchical grouping to optimize an objective function. Journal of the American Statistical Association 58(301), 236–244 (1963) 10.1080/01621459.1963.10500845

[30] Newman, M.E.J.: Networks: an Introduction, 1st edn. Oxford University Press, UK (2010)

[31] Zomorodian, A.: Fast construction of the Vietoris-Rips complex. Computers & Graphics 34(3), 263–271 (2010) 10.1016/j.cag.2010.03.007. Shape Modelling International (SMI) Conference 2010

[32] Puzis, R., Purohit, M., Subrahmanian, V.S.: Betweenness computation in the single graph representation of hypergraphs. Social Networks 35(4), 561–572 (2013) 10.1016/j.socnet.2013.07.006

[33] Takai, Y., Miyauchi, A., Ikeda, M., Yoshida, Y.: Hypergraph Clustering Based on PageRank. In: Proceedings of the 26th ACM SIGKDD International Conference on Knowledge Discovery & Data Mining. KDD ’20, pp. 1970–1978. Association for Computing Machinery, New York, NY, USA (2020). 10.1145/3394486.3403248

[34] Freeman, L.C.: A set of measures of centrality based on betweenness. Sociometry, 35–41 (1977) 10.2307/3033543

[35] Page, L., Brin, S., Motwani, R., Winograd, T.: The PageRank Citation Ranking: Bringing Order to the Web. Technical Report 1999-66, Stanford InfoLab (1999). http://ilpubs.stanford.edu:8090/422/

[36] Jaccard, P.: Distribution de la flore alpine dans le bassin des Dranses et dans quelques régions voisines. Bulletin de la Société Vaudoise des Sciences Naturelles 37(140), 241–272 (1901) 10.5169/seals-266440

[37] Kriege, N.M., Johansson, F.D., Morris, C.: A survey on graph kernels. Applied Network Science 5(1), 6 (2020) 10.1007/s41109-019-0195-3

[38] Vert, J.-P., Tsuda, K., Schölkopf, B.: A primer on kernel methods. Kernel Methods in Computational Biology, 35–70 (2004)

[39] Shawe-Taylor, J., Cristianini, N.: Kernel Methods for Pattern Analysis. Cambridge University Press, Cambridge (2004). 10.1017/CBO9780511809682

[40] Schölkopf, B., Smola, A.J.: Learning with Kernels: Support Vector Machines, Regularization, Optimization, and Beyond. The MIT Press, Cambridge (2002)

[41] Levenshtein, V.I.: Binary codes capable of correcting deletions, insertions, and reversals. Soviet physics doklady 10(8), 707–710 (1966)

[42] Yujian, L., Bo, L.: A Normalized Levenshtein Distance Metric. IEEE Transactions on Pattern Analysis and Machine Intelligence 29(6), 1091–1095 (2007) 10.1109/TPAMI.2007.1078

[43] Narayanan, A., Chandramohan, M., Venkatesan, R., Chen, L., Liu, Y., Jaiswal, S.: graph2vec: Learning distributed representations of graphs. CoRR abs/1707.05005 (2017)

[44] Le, Q., Mikolov, T.: Distributed representations of sentences and documents. In: Xing, E.P., Jebara, T. (eds.) Proceedings of the 31st International Conference on Machine Learning. Proceedings of Machine Learning Research, vol. 32, pp. 1188–1196. PMLR, Beijing, China (2014). https://proceedings.mlr.press/v32/le14.html

[45] Shervashidze, N., Schweitzer, P., Van Leeuwen, E.J., Mehlhorn, K., Borgwardt, K.M.: Weisfeiler-Lehman graph kernels. Journal of Machine Learning Research 12(9) (2011)

[46] Mikolov, T., Chen, K., Corrado, G., Dean, J.: Efficient estimation of word representations in vector space. arXiv preprint (2013) 1301.3781

[47] Hopkins, B.: A new method for determining the type of distribution of plant individuals. Annals of Botany 18(2), 213–227 (1954) 10.1093/oxfordjournals.aob.a083391. With an appendix by J. G. Skellam

[48] Banerjee, A., Dave, R.N.: Validating clusters using the Hopkins statistic. In: 2004 IEEE International Conference on Fuzzy Systems, vol. 1, pp. 149–1531 (2004). 10.1109/FUZZY.2004.1375706

[49] Cross, G.R., Jain, A.K.: Measurement of clustering tendency. IFAC Proceedings Volumes 15(1), 315–320 (1982) 10.1016/S1474-6670(17)63365-2. IFAC Symposium on Theory and Application of Digital Control, New Dehli, India, 5-7 January

[50] Kramer, M.A.: Nonlinear principal component analysis using autoassociative neural networks. AIChE Journal 37(2), 233–243 (1991) 10.1002/aic.690370209

[51] Vincent, P., Larochelle, H., Bengio, Y., Manzagol, P.-A.: Extracting and composing robust features with denoising autoencoders. In: Proceedings of the 25th International Conference on Machine Learning, pp. 1096–1103 (2008)

[52] Bai, S., Zhang, F., Torr, P.H.: Hypergraph convolution and hypergraph attention. Pattern Recognition 110, 107637 (2021) 10.1016/j.patcog.2020.107637

[53] Pękalska, E., Duin, R.P.W.: The dissimilarity representation for pattern recognition: foundations and applications. Series in Machine Perception and Artificial Intelligence. World Scientific (2005). 10.1142/5965

[54] De Santis, E., Martino, A., Rizzi, A.: On component-wise dissimilarity measures and metric properties in pattern recognition. PeerJ Computer Science 8, 1106 (2022) 10.7717/peerj-cs.1106

[55] Pękalska, E., Paclik, P., Duin, R.P.W.: A generalized kernel approach to dissimilarity-based classification. J. Mach. Learn. Res. 2, 175–211 (2002)

[56] Martino, A., Giuliani, A., Rizzi, A.: (Hyper)Graph Embedding and Classification via Simplicial Complexes. Algorithms 12(11) (2019) 10.3390/a12110223

[57] Rosenberg, A., Hirschberg, J.: V-measure: A conditional entropy-based external cluster evaluation measure. In: Proceedings of the 2007 Joint Conference on Empirical Methods in Natural Language Processing and Computational Natural Language Learning (EMNLP-CoNLL), pp. 410–420 (2007)

[58] Manning, C.D., Raghavan, P., Schütze, H.: Flat clustering. In: Introduction to Information Retrieval, pp. 349–375. Cambridge University Press, Cambridge (2008)

[59] Rousseeuw, P.J.: Silhouettes: a graphical aid to the interpretation and validation of cluster analysis. Journal of Computational and Applied Mathematics 20, 53–65 (1987) 10.1016/0377-0427(87)90125-7

[60] McInnes, L., Healy, J., Saul, N., Großberger, L.: Umap: Uniform manifold approximation and projection. Journal of Open Source Software 3(29), 861 (2018) 10.21105/joss.00861

[61] McInnes, L., Healy, J., Melville, J.: UMAP: Uniform Manifold Approximation and Projection for Dimension Reduction (2020). https://arxiv.org/abs/1802.03426

